# Ovo, an Open-Source Ecosystem for *De Novo* Protein Design

**DOI:** 10.1101/2025.11.27.691041

**Authors:** David Prihoda, Marco Ancona, Tereza Calounova, Adam Kral, Lukas Polak, Hugo Hrban, Nicholas J. Dickens, Danny A. Bitton

**Affiliations:** Applied Research and Innovation, MSD Czech Republic s.r.o., Prague, Czech Republic; Amazon Web Services LLC, Seattle, WA, United States

## Abstract

The protein design field is rapidly advancing, with frequent emergence of new models and pipelines for designing de novo proteins with tailored properties and functions not found in nature. However, the current tool landscape is fragmented, tools are hard to install and deploy, and require significant computational expertise to integrate into end-to-end, scalable pipelines. A particular challenge is managing many sequences, structures, and metrics for downstream testing and retrospective analysis of input parameters. To address this need, we introduce Ovo, an open-source de novo protein design ecosystem that consolidates models, workflows, data management, and interactive visualization into a scalable, infrastructure-agnostic platform. Ovo features Nextflow-based workflow orchestration, a storage layer, and both command-line and graphical interfaces that democratize scaffold design, binder design and diversification, and validation workflows. Ovo’s novel ProteinQC module computes comprehensive sequence and structure descriptors, contextualizing designs against reference sets. Ovo plugins let the community add new workflows and user interfaces to accelerate adoption of emerging methods and facilitate community-driven benchmarking. Ovo lowers engineering barriers and demystifies the design process, allowing experts and non-technical users to design proteins at scale. With community-driven development, Ovo can accelerate de novo protein design and advance discovery in therapeutics and biotechnology.

## Introduction

Recent breakthroughs in computational biology have sparked an outburst of development of protein folding^1–4^, inverse folding^5,6^, co-folding^7–10^and generative diffusion models^11–13^, fundamentally transforming protein design, with profound implications for drug and vaccine development and manufacturing^11,14^.

These advancements have dramatically expanded the protein engineering toolbox, enabling accurate prediction of a three-dimensional protein structure from sequence^1–4,7–10^, generation of sequences that can fold into a given three-dimensional backbone^5,6,15,16^, modeling of protein-protein^2,7–10^ and protein-ligand interactions^7–10,16,17^ and even generating entirely novel protein scaffolds and binders that do not exist in nature^11,13,15,18^. The latter typically involves chaining multiple models in sequence to create powerful *in silico* design pipelines^11,15,18,19^. These *in silico* pipelines can generate thousands of protein backbones and sequences with desirable properties and functions based on defined input criteria. Thereafter, the resulting designs are rigorously filtered using various confidence measures and scores from independent models, yielding a handful of carefully selected candidates that are further subjected to experimental validation via downstream binding and functional assays. An increasing number of studies have demonstrated successful *de novo* design of functional proteins^11,15^, enzymes^13^, mini-protein binders^11,14,18–20^, antibodies^21–23^, short linear and cyclic peptides^18,24–26^with a broad applicability in medicine and biotechnology.

However, navigating the vast sea of protein modelling tools is far from trivial, let alone establishing robust and scalable pipelines for *de novo* protein design that can meet industrial throughput needs and standards. These models and pipelines are sparsely distributed, some as standalone command line tools, notebooks or scripts, often lacking standardized installation instructions or comprehensive documentation. Setting up these tools can be cumbersome, requiring understanding of numerous model parameters and settings as well as significant computational expertise to setup the compute resources and to manage software dependencies and versions. Moreover, complexity increases with the need to integrate multiple tools into cohesive pipelines, where the output of one model serves as the input for another. This requires careful management of settings, data formats, and a tight orchestration of multiple processes, further complicating reproducibility and scalability. Furthermore, because the field is still in its infancy, new models are rapidly emerging, with variable reports on experimental success, creating a real need to rapidly test, adopt and integrate top-performing models.

Recently, Nextflow^27^ and the nf-core^28^ community streamlined and simplified workflow development by using a flexible workflow language and containerized processes to improve modularity, scalability and reproducibility. Nextflow provides abstraction of backend execution, and easy integration of containers, enabling parallelism and execution on a single machine, High Performance Computing (HPC), or cloud infrastructure. The nf-core community provides community-curated pipeline templates, automated testing and continuous integration as well as comprehensive documentation and guidelines to ensure portability, testing, and best practices. Together, these initiatives reduce setup effort, improve reproducibility, and accelerate the sharing and reuse of robust bioinformatics workflows by the broader community.

Even after pipelines are established, substantial challenges remain. An intuitive user interface is needed to help non-technical users choose appropriate models and workflows for a given design task (i.e. scaffold design, binder design or diversification) and invoke pipelines within large-scale compute infrastructure. Upon retrieval of thousands of designs, the predicted structures and their associated confidence scores are often insufficient to identify the best candidates. Scientists frequently require additional features, sequence-, structure-, and physics-based information to make data-driven decisions. Thus, users must interactively evaluate and visually inspect these candidates, apply varying thresholds of filters based on confidence and pre-computed design properties, and ultimately select most promising designs for experimental validation.

Moreover, in pharmaceutical environments *de novo* drug design campaigns often produce hundreds of thousands of designs across many targets and pipelines. These campaigns are dynamic and collaborative and may be subject to future regulatory scrutiny. Therefore, to ensure reproducibility, metadata, pipelines, and results must be versioned, recorded, and fully documented, which requires robust data management capabilities.

Here we introduce Ovo, an open-source *de novo* protein design ecosystem tailored to address these needs. Ovo offers an intuitive and interactive visualization and workflow orchestration platform for an end-to-end *de novo* protein design that can be readily deployed on the institution’s HPC or cloud infrastructure. Ovo is powered by a robust, scalable, high-performance, modular and extensible backend foundation that features key models, pre-defined workflows, and advanced filtering capabilities based on confidence scores and numerous protein descriptors. Ovo not only provides a unified framework that demystifies the *de novo* design process but also aims to empower the community to evaluate new methods, standardize workflows, and develop additional pipelines and plugins that further advance the protein design field.

## Results

### Ovo - an intuitive, user-friendly, scalable and extensible de novo design ecosystem

Ovo aims to streamline and simplify *de novo* protein design. To this end, the following principles guided its planning and development. A key goal was to encompass all common *de novo* protein design workflows, including scaffold design, binder design and diversification and co-folding. On the technical front, we aimed to deliver a plug-and-play, extensible ecosystem in which different models and filters can be mixed and matched to create bespoke pipelines. A critical prerequisite was an infrastructure-agnostic platform that can be readily deployed on local machines, HPC, or in the cloud. Given the anticipated large number of design campaigns run by multiple concurrent users in pharmaceutical settings, the system must offer scalable, parallel computation to minimize runtime. Moreover, the system should serve power users by offering programmatic access and advanced configurable settings via a command-line interface (CLI), while also enabling non-technical modelers to invoke scalable pipelines via an interactive user interface (UI). The development of a data management layer was also a key requirement to ensure efficient organization, storage, and retrieval of designs and descriptors across multiple concurrent projects and users. The data model should facilitate retrospective analysis, relating experimental success rates with parameter choices and computational scores through unique identifiers. In addition, the software must provide an advanced, interactive, and intuitive visualization layer that allows filtering of resulting designs based on confidence measures and protein characterization features. Lastly, exporting designs and integrating with Jupyter notebooks and third-party software for downstream analysis should be seamless and straightforward.

Architecturally, Ovo is comprised of four main components: web application (Ovo app), command-line interface (Ovo CLI), logical layer (Ovo core) and Nextflow pre-defined workflows (Ovo pipelines, Figure 1a). With the Ovo CLI and Ovo app, both technical and non-technical users can submit predefined computational workflows at scale, then visually inspect and filter resulting designs using confidence scores and protein properties filters (Figure 1b). As a Python library, Ovo also provides a programmatic interface for seamless integration with Jupyter notebooks and third-party tools, enabling downstream analysis and further refinement of designs. Ovo workflows each chain a configurable selection of models to enable scaffold design, binder design and diversification and protein characterization, as discussed in detail in subsequent sections (Figure 1c, Table 1). Community developed workflows and tools can be readily integrated to Ovo as plugins (Figure 1d).

**Figure 1.**
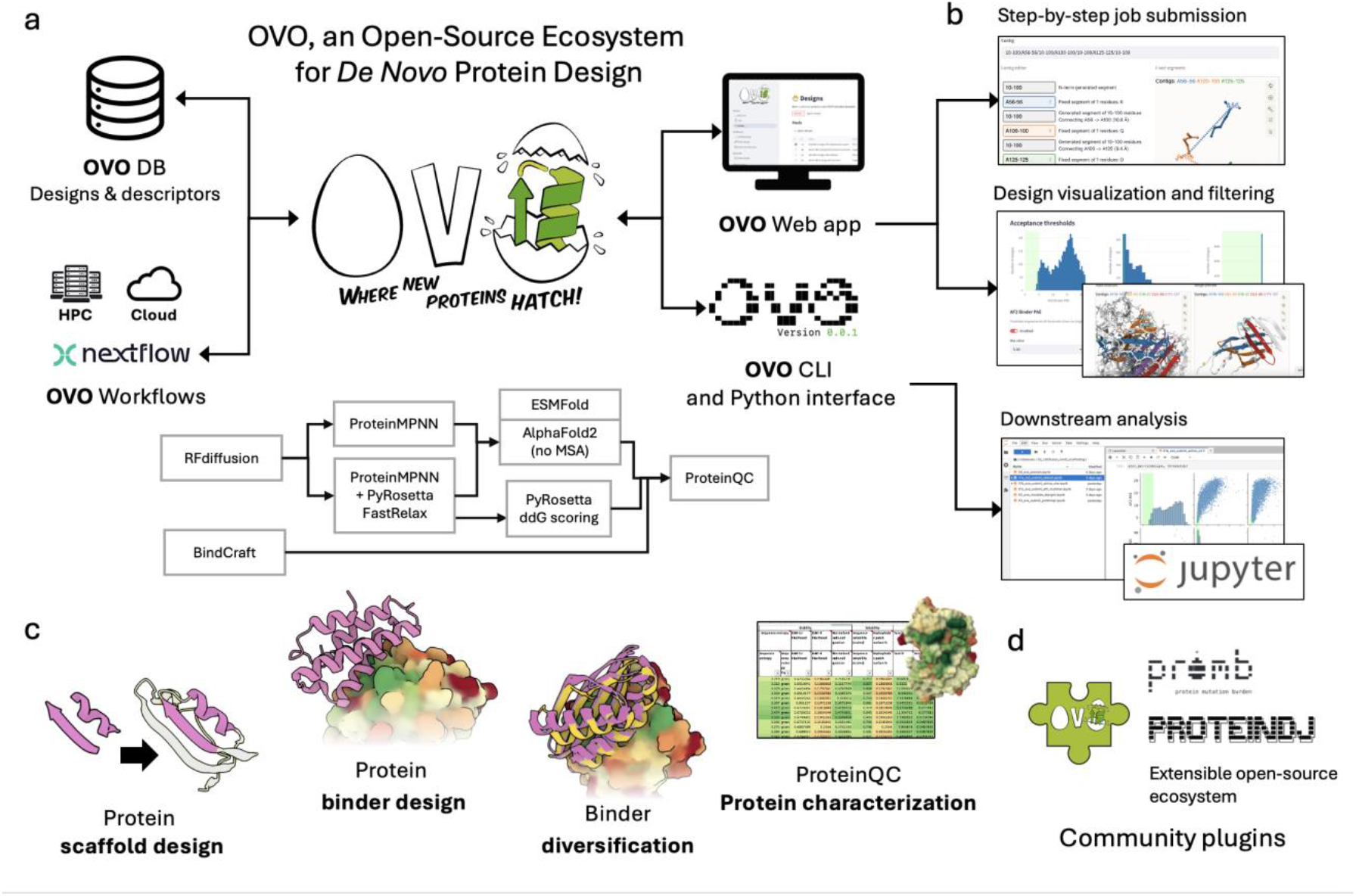
An overview of Ovo. (a) A high-level architecture diagram showing the main components: an SQL database to store designs, descriptors, and their metadata in a standardized data model; a workflow definition and orchestration system based on Nextflow; a web application for step-by-step workflow submission, design visualization and filtering; and a command-line interface or Python interface for programmatic access and seamless downstream analysis in Jupyter notebooks or third-party software. (b) a closer look at the Ovo interface that allows user-friendly workflow submission, interactive visualization and filtering of resulting designs based on various confidence scores and filters (c) Pre-defined workflows in Ovo for design of scaffolds (RFdiffusion, ProteinMPNN, AlphaFold2), binders (RFdiffusion and BindCraft), binder diversification (RFdiffusion) and ProteinQC protein characterization. (d) Ovo supports integration with community developed tools and workflows as demonstrated by plugins for ProteinDJ protein design pipeline or Promb protein humanness evaluation.

**Table 1.**
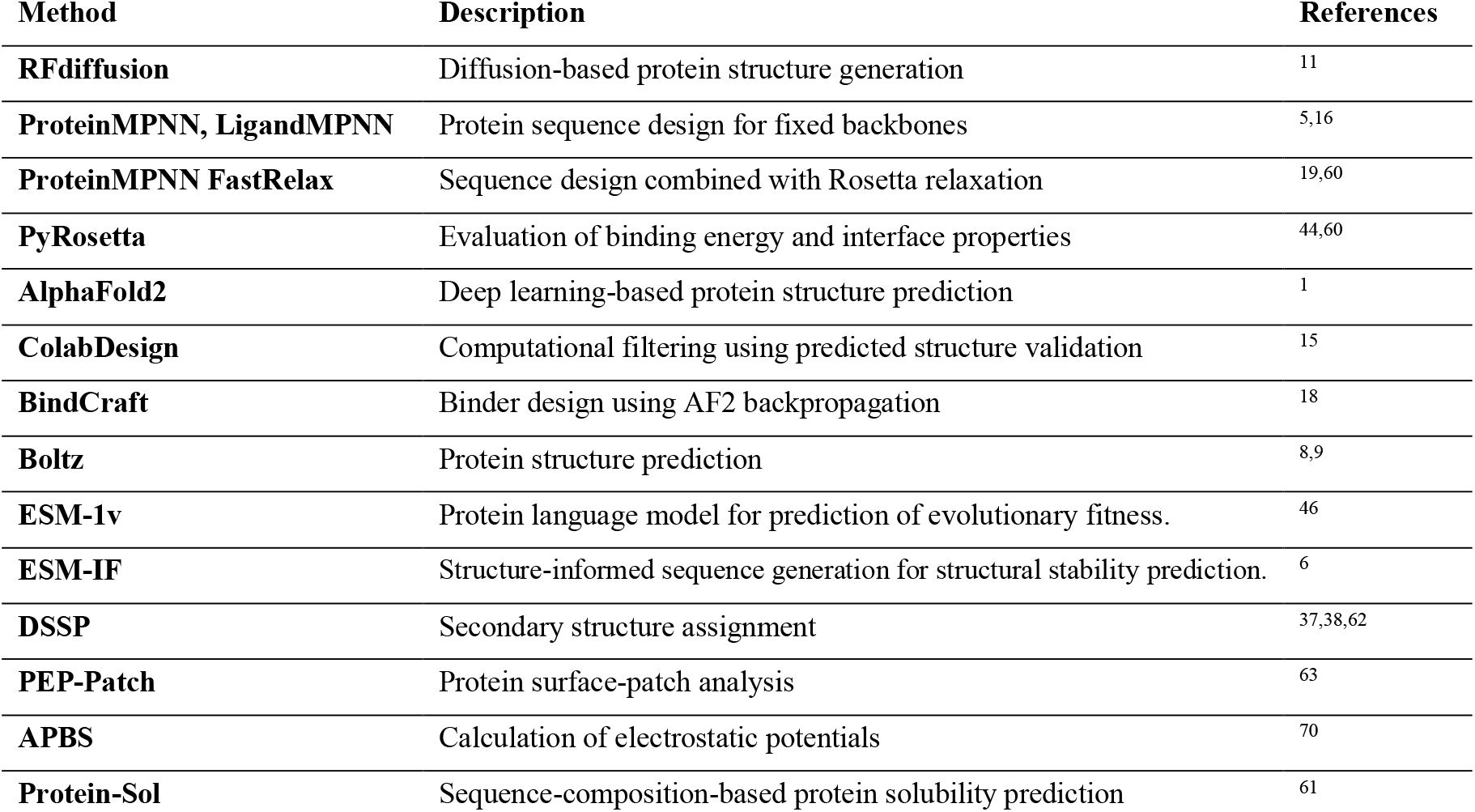
The tools and models that are integrated into Ovo and enable *de novo* design of protein scaffold and binders design and diversification.

Taken together, Ovo is a unified, extensible platform that combines scalable, infrastructure-agnostic computation with an intuitive interface and programmatic access, empowering both expert and non-technical users to run end-to-end *de novo* protein design workflows.

### Ovo enables functional motif scaffolding around enzyme active sites and binding interfaces

*De novo* scaffold design refers to the generation of an entirely new protein backbone that supports an existing functional site (motif) of a protein. It has a wide range of applications in therapeutics, drug manufacturing, and diagnostics. For example, scaffold design workflows can be used to create epitope-focused vaccines^29,30^, amplify immune responses by constructing multivalent antigen structures^11,31–34^ or design compact scaffolds around an enzyme’s active-site geometry to improve stability or solubility^35^. Similarly, these workflows can be applied to miniaturize a known protein binder, preserving only its interface in a new stable scaffold^11^.

Ovo offers an end-to-end functional motif scaffolding workflow that can generate new backbones that preserve user-defined fixed segments and produces compatible sequences and side-chain conformations. Ovo’s scaffold design workflow integrates RFdiffusion^11^ for *de novo* backbone generation, LigandMPNN^16^ (with ProteinMPNN^5^ model weights) for sequence and rotamer prediction, and AlphaFold2 (AF2)^1^ and/or ESMFold^4^ for orthogonal structure prediction and filtering.

Starting from a source structure PDB file or PDB ID, users select fixed residue segments and interactively specify a “contig” string that defines the order in which the fixed stretches should be connected (original order by default, but alternative permutations can be considered) and how many residues to design between them, and on N/C termini. Users can verify the contig by generating a backbone preview using a reduced number of RFdiffusion timesteps (T=1-20). Next, users can interactively select residues for sequence inpainting, which are positions where the backbone is fixed but the sequence is hidden from RFdiffusion. These residues are then redesigned using ProteinMPNN, enabling improved packing. The submission form allows users to specify the numbers of backbones and sequences along with all advanced parameters for RFdiffusion, ProteinMPNN and AlphaFold2. This workflow is modular and scalable but computationally demanding. Robust RFdiffusion campaigns typically require many thousands of backbone generations to recover a small fraction of high-confidence designs, so users are advised to run an initial pilot of 100–200 backbones to validate inputs and settings before scaling up. Upon workflow completion, resulting designs are registered in the Ovo database and presented in a dedicated “Job Results” page in the UI, where users can visualize structure alignments, review design descriptors, adjust selection thresholds, and select top designs (typically less than ∼0.1% of generated sequences) for further analysis in a designated “Designs” page. Designs that pass the acceptance thresholds can be further analyzed with descriptor workflows such as ProteinQC, which can be triggered from the interface to compute a complete set of protein-characterization descriptors (Supplementary Table S1) as described in the designated ProteinQC section. Figure 2 illustrates scaffold designs around an input functional motif of Oxidoreductase (Figure 2a-c), and around the natural binding interface of PD-1 (Figure 2d-e) generated by Ovo based on previous reports^11,36^. Workflow parameters and summary statistics are provided in the Methods section and in Supplementary Tables S2 and S3.

**Figure 2.**
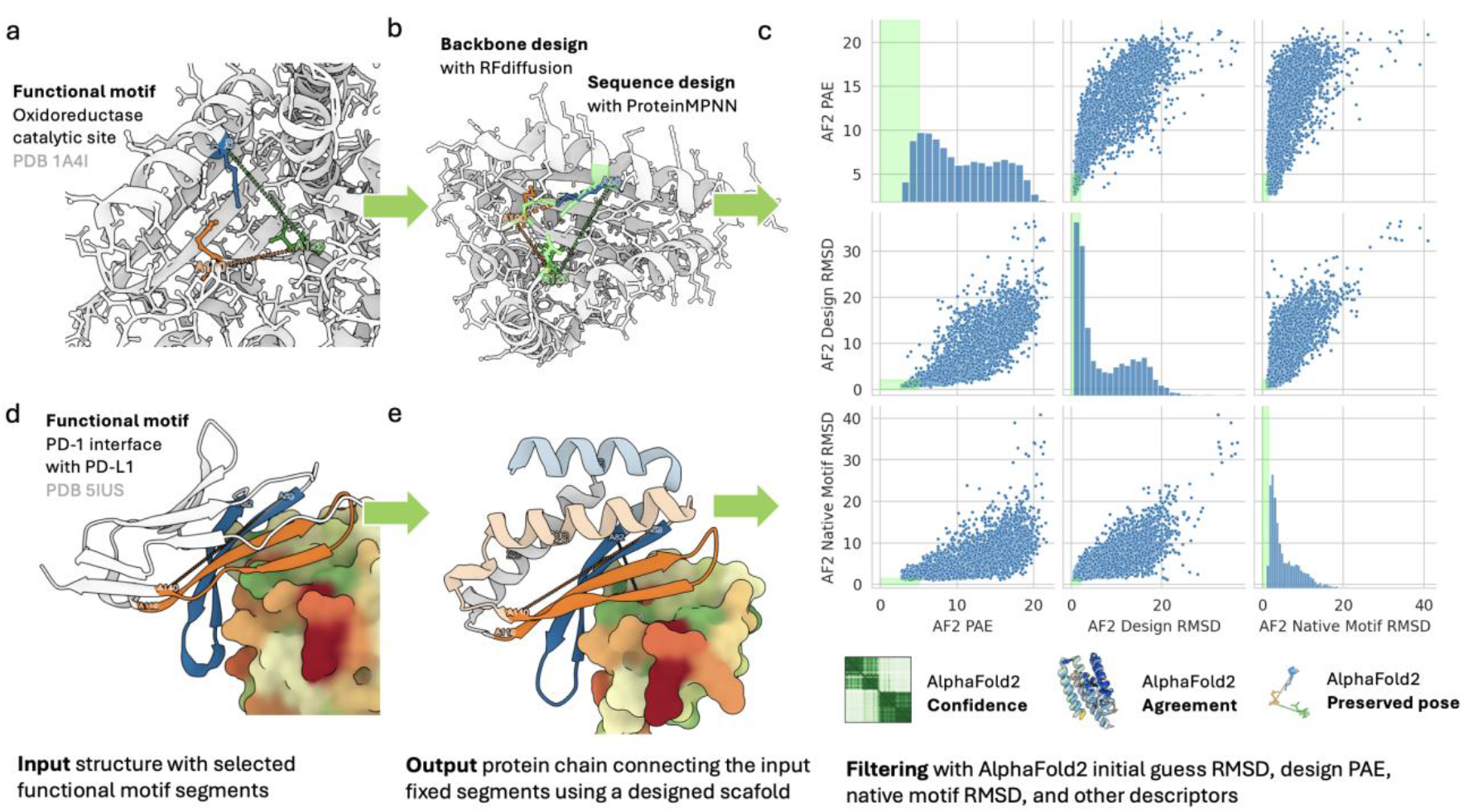
Functional motif scaffolding using Ovo (a-b) example design of a scaffold around input motif of three active-site residues of oxidoreductase (PDB accession: 1A4I)^66^ in a similar fashion to the one designed by Watson *et al*.^11^ (c) Filtering using AlphaFold2 predicted structure validation metrics demonstrated on oxidoreductase designs: mean PAE < 5 Å, mean pLDDT > 80, Design RMSD < 2 Å and Native Motif RMSD < 1.5 Å computed using 3 recycles of AlphaFold2 initial guess monomer model 1 with fixed input motifs used as structure templates. (d-e) example design of a scaffold (grey, light orange and light blue helices) around the interface of PD-1 (orange and blue, PDB accession 5IUS)^67^ binding PD-L1 (surface representation colored by hydrophobicity, hydrophobic = green, hydrophilic = red) following the same design principle reported in Wang *et al*.^36^, core-facing residues masked with RFdiffusion sequence inpainting and redesigned with ProteinMPNN.

Importantly, output designs are accompanied by backbone, sequence-composition, and predicted structure validation (“refolding”) descriptors that together provide a compact, independent assessment of whether an RFdiffusion design is structurally plausible, chemically sensible, and likely to fold into the intended conformation. The complete set of descriptors computed by Ovo, together with brief descriptions, comparison direction (higher vs. lower preferred), value ranges, and warning thresholds, is provided in Supplementary Table S1. Ovo computes backbone metrics with PyDSSP^37,38^ and custom logic, reporting percentages of helix, sheet, and loop and the radius of gyration. These metrics evaluate the geometric plausibility and compactness of the fold, which can in turn affect function, solubility, and stability. Similarly, Ovo computes sequence-composition descriptors using the Biopython ProtParam (ProteinAnalysis) module^39^ to flag biochemical and manufacturability risks. Reported features include amino acid composition, net charge at pH 7.4 and 5.5, isoelectric point (pI), sequence entropy, and aromaticity. These metrics help users assess factors that may affect expression, aggregation, solubility, and stability of the newly designed scaffolds.

Ovo also computes predicted structure validation metrics using AlphaFold2 initial guess^19^ or ESMFold that predict whether the designed sequence is likely to fold into the intended structure and preserve the user-specified functional motif. Such metrics include AlphaFold2’s Predicted Aligned Error (mean PAE) as a global confidence estimate, the Design RMSD (Cα Root Mean Square Deviation between the RFdiffusion backbone and the AlphaFold2 prediction) to assess whether the ProteinMPNN-designed sequence is likely to fold into the intended backbone. RFdiffusion does not preserve side-chain atoms and does not guarantee the preservation of the positions of fixed backbone atoms, especially in complicated contigs with many separate fixed segments. Therefore, the Native Motif RMSD (RMSD between all atoms of the functional motif in the input structure compared to the predicted structure) is computed to confirm preservation of the functional motif. Designs are marked as accepted with default thresholds of mean PAE < 5 Å, mean pLDDT > 80, Design RMSD < 2 Å and Native Motif RMSD < 2 Å, computed using 3 recycles of AlphaFold2 initial guess monomer model 1 with fixed input motifs used as structure templates. These default thresholds can be adjusted by the user before or after workflow submission. Once filtering reduces the design pool from tens of thousands to a few hundred accepted candidate designs, users are encouraged to use the Refolding workflow to submit additional validation using different models, settings, and apply more stringent thresholds. Per-residue and global confidence scores such as pLDDT (predicted Local Distance Difference Test) and pTM (predicted Template Modelling) are also reported to assess local reliability and overall fold quality, respectively. An example of filtering thousands of designs based on AF2 RMSD and PAE is shown in Figure 2c.

We conclude that Ovo streamlines the design of bespoke protein backbones that preserve functional sites. Its interactive UI lets users launch scalable pipelines and prioritize designs by structural, sequence, and refolding metrics, enabling rational selection of high-quality candidates for downstream experimental validation.

### Ovo features interactive and modular binder design capabilities

*De novo* biological binder design refers to the generation of entirely new proteins or short peptides that are engineered to form specific interactions with a defined binding site on a given molecular target even in the absence of a natural binder. Biological binders form the foundation of modern therapeutics, as they can bind specific targets such as receptors, enzymes, or signaling molecules and thereby modulate cellular pathways to achieve a desirable therapeutic effect. Their capacity to control biological functions with high specificity makes biological binders such as antibodies, proteins, and peptides attractive candidates for engineering. Such engineering efforts typically aim to improve specificity, stability, immunogenicity profile, and ADMET properties (absorption, distribution, metabolism, excretion, toxicity), and make these molecules easier and less expensive to manufacture at scale. In this context, mini-proteins and short peptides offer several advantages over larger molecules such as antibodies, including improved drug-like properties, simpler manufacturability, and greater amenability to *de novo* design. Yet their application extends beyond therapeutics to diagnostic^24,40,41^, biochemical assays^42^, and manufacturing as affinity reagents^43^.

Ovo provides bespoke end-to-end mini-protein and linear peptide design pipelines that generate binders to a user-specified target surface, featuring two literature-validated protocols based on RFdiffusion^11^ and BindCraft^18^. Users begin by providing a target structure and selecting the target chain. Optionally, they can specify trimming boundaries to reduce computation time and GPU memory requirements. Users must also define the binder chain length range and may optionally list hotspot residues to bias the binding site (Figure 3a), along with any additional advanced parameters (e.g. filtering criteria, enriching or omitting specific amino acids etc.). As noted previously, Ovo offers a quick preview mode to validate inputs prior to triggering large-scale runs. It also records all output designs and descriptors in Ovo’s database and enables interactive inspection and filtering via the UI.

**Figure 3.**
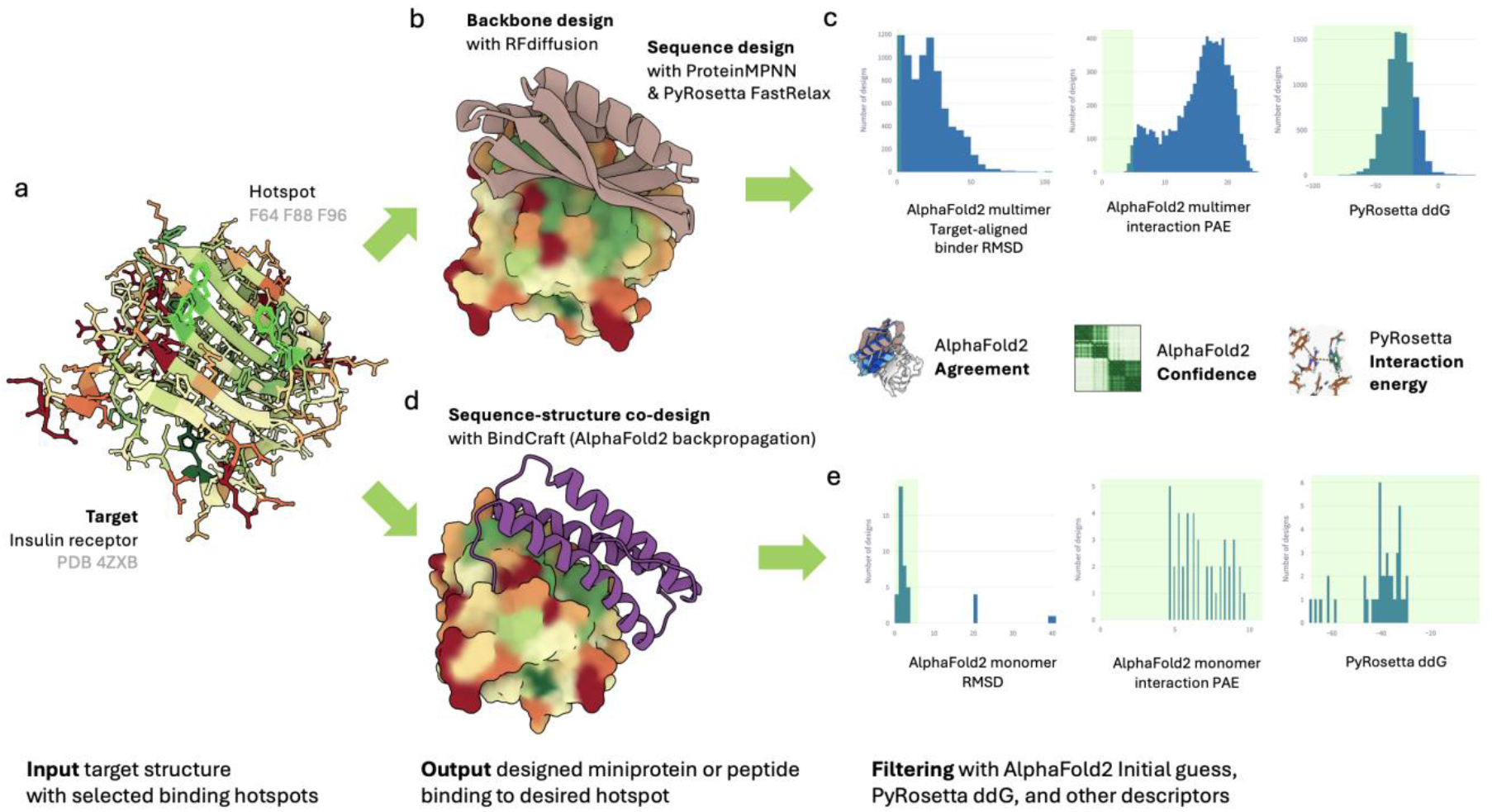
Binder design with Ovo (a) Insulin receptor input structure in Ovo’s user interface (PDB accession 6PXV) ^68^ colored by hydrophobicity (green = hydrophobic, red = hydrophilic) with selected hotspot shown with a green outline (b) An example of a binder designed with the RFdiffusion workflow followed by filtering with AlphaFold2 initial guess. RFdiffusion generates a backbone, while PyRosetta FastRelax and ProteinMPNN protocol generates sequences and side-chain rotamers. (c) AlphaFold2 evaluates design by re-predicting the complex, computing the confidence of the prediction, and by comparing the binder pose from the RFdiffusion backbone to the AlphaFold2 prediction (d) Binder design to the same insulin receptor structure using BindCraft workflow. BindCraft uses AlphaFold2 multimer backpropagation, followed by ProteinMPNN surface sequence design, PyRosetta scoring and various filters including (e) refolding metrics that are derived from AlphaFold2 monomer and interface metrics computed with PyRosetta.

#### Binder design with RFdiffusion

The RFdiffusion workflow generates *de novo* binder backbones in the context of the trimmed target. Sequences and side-chain rotamers are produced by iterative ProteinMPNN design coupled with PyRosetta FastRelax^19^, or by LigandMPNN^16^ (with ProteinMPNN^5^ model weights). Thereafter, designs are evaluated by AlphaFold2 multimer initial guess, producing the confidence measures.

Similar to the evaluation of *de novo* scaffolds, Ovo computes backbone, sequence-composition and predicted structure validation metrics to enable systematic filtering of the generated designs with notable differences described below. On top of the secondary-structure descriptors computed by PyDSSP and custom functions, Ovo also calculates backbone–backbone residue proximity counts between binder and target hotspots. This measures how extensively the binder and target interact and specifically whether the binder engages the desired epitope (since RFdiffusion does not guarantee this placement). Greater total backbone contacts and more contacts with hotspot residues indicate a larger, more shape-complementary interface that can stabilize and enhance affinity, while being more tolerant to side-chain changes. Designs with low backbone-hotspot engagement may rely on precise side-chain packing and be less robust, so these metrics may help prioritize robust, high-affinity candidates. Backbones that do not satisfy user-defined thresholds can be discarded after backbone generation, saving compute time and resources. In addition to sequence-composition descriptors described earlier (amino-acid frequencies, net charge at pH 7.4/5.5, pI, entropy and aromaticity), Ovo also computes interface descriptors using PyRosetta^44^. The Rosetta ddG predicts binding energy, with more negative values indicating stronger predicted affinity. The contact molecular surface measures the buried surface area at the interface where larger values indicate a bigger contact patch and often stronger or more specific interactions. The spatial aggregation propensity (SAP) score evaluates exposed hydrophobicity, positive scores implying a greater aggregation risk. Therefore, by combining these interface metrics, users may select designs with strong predicted binding, substantial interfaces, and low aggregation propensity.

Lastly, each design is independently validated by AlphaFold2 initial guess structure prediction^19^ to identify those with consistent and confident binder–target poses. AlphaFold2 offers per-residue and global confidence scores (local pLDDT and global pLDDT or pTM, respectively), pairwise confidence (predicted aligned error, PAE), and confidence of the interface (interaction PAE, or sometimes called interface PAE, abbreviated as iPAE, or ipAE) corresponding to the predicted error of binder–target pose in angstroms (Å), computed as the mean over all binder–target residue pairs in the PAE matrix. The Binder RMSD is the Cα RMSD of the binder between the RFdiffusion design and the AlphaFold2 prediction after alignment on the target, evaluating agreement with respect to the binder’s backbone pose and placement on the target. Designs are marked as accepted with default thresholds of iPAE < 10, pLDDT > 80, and Target-aligned Binder RMSD < 2 Å, computed using AlphaFold2 initial guess multimer model 1 with target chain used as structure template, and Rosetta ddG < −30. Once filtering reduces the design pool from tens of thousands to a few hundred accepted candidate designs, users are encouraged to use the Refolding workflow to submit additional validation using different models, settings, and apply more stringent thresholds. RFdiffusion typically requires large sampling, often tens of thousands of backbones to recover a small fraction of high-confidence designs. Since RFdiffusion workflows are computationally intensive, a pilot run of 100–200 backbones is typically recommended prior to launching large-scale jobs.

#### Binder design with BindCraft

BindCraft^18^ is a three-step workflow that uses AlphaFold2 backpropagation via ColabDesign^15^ to iteratively hallucinate binder sequences and poses from randomly initialized starting points, followed by surface sequence redesign with SolubleMPNN^45^ or ProteinMPNN, PyRosetta scoring, and advanced filtering. First, it iteratively hallucinates binder sequences and poses using backpropagation through AlphaFold2 multimer network to optimize confidence metrics, interchain contacts, and other differentiable objectives. Thereafter, it performs a sequence redesign that resamples noninterface positions with SolubleMPNN (default) or ProteinMPNN to improve expression and solubility^18^. The third stage is the filtering phase, where the workflow uses Alphafold2 monomer prediction as an orthogonal check that the interface is well defined even according to a model that has never been exposed to multimeric complexes. Multiple filters are then applied to the designs by default, including AlphaFold2 confidence metrics (ipTM > 0.5 and normalized iPAE < 0.35, equivalent to < 10.85 in Å) and low AlphaFold2 Hotspot RMSD (< 6 Å) that ensures compatibility between the original hallucinated binder pose with AlphaFold2 multimer with the MPNN-redesigned AlphaFold2-monomer reproduced binder. In addition, unstructured binders, and binders that do not meet interface residue contact thresholds are filtered out by default. Similar to the RFdiffusion workflow, accepted BindCraft designs include binder sequences alongside predicted complex structures and design descriptors, for example predicted refolding metrics, PyRosetta scores such as ddG, buried contact surface and SAP, and secondary structure annotations. Compared with RFdiffusion, BindCraft often yields more accepted designs per unit compute but is more GPU memory intensive since backpropagation through AlphaFold2 requires storing all intermediate layers’ activations unlike a forward pass through a diffusion model. Users encountering out-of-memory errors therefore should trim the target or alternatively use larger GPU instances.

Once the RFdiffusion or BindCraft workflows have finished, all generated designs and their descriptors are saved to Ovo storage and registered in the Ovo database so users can view them in the “Job Results” page. For RFdiffusion, users can adjust the acceptance criteria after workflow completion. Typically, only ∼0.1%–1% of RFdiffusion outputs are accepted. BindCraft designs are rejected or accepted directly as part of the workflow based on the filters configured at submission time. Designs marked as accepted are available for further analysis and filtering in the “Designs” page by descriptor workflows such as ProteinQC. Examples of binder design to the insulin receptor by Ovo’s RFdiffusion and BindCraft workflows are shown in Figure 3a-c and Figure 3a, d and e, respectively. Workflow parameters and result statistics are provided in the Methods section and in Supplementary Tables S2 and S3.

In summary, Ovo integrates RFdiffusion and BindCraft into modular, parallelized binder design workflows that couple large-scale backbone sampling and sequence design with independent AlphaFold2 validation to prioritize structurally and chemically plausible binders for experimental follow-up.

### Ovo features binder diversification with partial diffusion

Binder diversification refers to the generation of a variety of binder backbones and sequences based on an existing binder template in the context of a binder-target complex. Typical applications of binder diversification include improving the affinity of an experimentally derived or computationally designed binder^14^, altering specificity by adapting the binder to a homologous target^14,20^, and increasing success rates by sampling around previously successful designs^20^.

Ovo offers a linear peptide and mini-protein binder diversification workflow that uses a partial RFdiffusion protocol that first adds noise to the binder backbone and then iteratively removes it to generate a similar backbone. Starting with one or more PDB structures (or Ovo’s internal design IDs) containing the binder and target chains, users can select to redesign the whole binder chain (default) or restrict the partial diffusion and subsequent sequence redesign to specific residues or segments of the binder. Compared to the binder interface scaffolding task described above, partial diffusion is only trained for fixed-length redesign where each residue in the input binder corresponds directly to a residue in the output, so the overall length and segment order are preserved. Users can also optionally specify hotspot residues to bias the binding towards a specific part of the target molecule and can adjust the noising strength to control the number of RFdiffusion iterations (default=20). Similar to the binder design workflow, Ovo’s diversification workflow outputs binder sequences and complex structures together with descriptors that include backbone metrics (secondary structure composition, backbone–backbone and hotspot contact counts), sequence composition (residue frequencies, charge at pH 7.4/5.5, pI, aromaticity), PyRosetta scores (ddG, contact molecular surface, spatial aggregation propensity) and AlphaFold2 validation metrics (iPAE, Target-aligned Binder RMSD, Binder PAE, Binder pLDDT and pTM). A full and restricted backbone diversification of a *de novo* designed candidate binder to the insulin receptor (PDB ID 6PXV) are illustrated in Figure 4a and b, respectively, followed by AF2 Binder RMSD, AF2 interaction PAE and design Rosetta ddG filtering (Figure 4c). Accepted designs along with their descriptors are stored in Ovo’s storage layer and can be further analyzed by ProteinQC workflow via the Ovo app.

Ovo’s diversification workflow can generate diverse binder backbones around an input binder–target complex using a partial RFdiffusion protocol, accelerating identification of binders with improved properties for downstream validation experiments.

**Figure 4.**
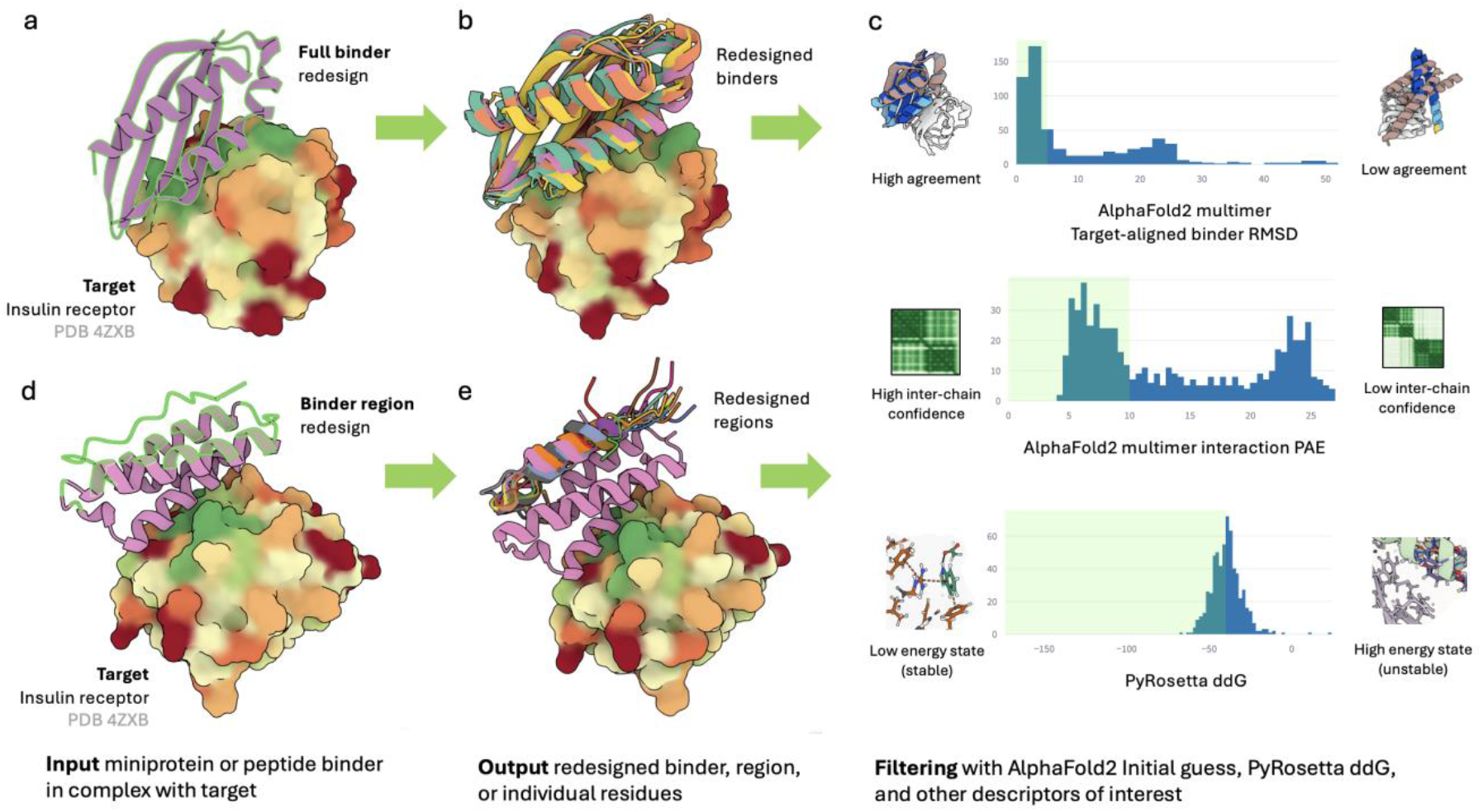
Binder diversification via Ovo’s partial RFdiffusion workflow. (a) A binder-target complex structure was used as an input (insulin receptor: PDB accession 6PXV and an accepted binder from Figure 3) and subjected to partial RFdiffusion using default settings (T=20 iterations). (b) showing 3 examples of accepted backbones that were evaluated using backbone and sequence-composition metrics, alongside PyRosetta scores and (c) AlphaFold2 validation metrics (acceptance criteria AF2 Interaction PAE < 10 Å, AF2 target-aligned binder RMSD < 2 Å and Rosetta ddG < −40). (d) Partial diffusion of non-interface residues, residues to be redesigned highlighted with a green overlay (e) Example diversification result showing multiple versions of a redesigned region of an existing binder design.

### Protein QC - evaluating designs with respect to features of a reference set

Ovo’s ProteinQC workflow is an automated, data-driven quality-control module that characterizes natural or *de novo* protein designs in the context of a reference set, helping to flag problematic chemistries, folding risks, stability issues, and expression or aggregation liabilities. The ProteinQC workflow can be triggered directly from the UI or the CLI to generate a complete set of descriptors (Supplementary Table S1) for accepted designs at the end of each design workflow, or for any user-uploaded pool of structures. These descriptors enable comparative analysis against a reference set (all PDB structures or specific PDB subsets), helping users interactively contextualize designs and export them as advanced, well-formatted Excel spreadsheets for downstream clustering and filtering. QC descriptors can be broadly divided into three distinct yet complementary classes that reflect their insight into stability, expression and solubility, and protein structure, as described below.

#### Stability descriptors reveal lower sequence entropy of de novo designs

Exemplary descriptors such as sequence entropy, ESM-1v^46^ (Evolutionary Scale Modeling) likelihood, and ESM-IF^6^ (ESM-Inverse Folding) likelihood are used by the ProteinQC workflow as sequence- and structure-derived proxies for stability, among many additional descriptors (Supplementary Table S1). Sequence entropy is computed as Shannon entropy of the amino acid distribution over a sliding window, reflecting local amino acid composition diversity, with low entropy indicating repetitive or low-complexity regions and vice versa. Repetitive regions are not necessarily a cause for concern but may imply higher risk of nonspecific interactions or misfolding, so they warrant further inspection. The ESM-1v protein language model is used to compute an evolutionary likelihood score, higher values indicating that the sequence conforms to patterns seen in natural protein sequences. This provides a faster alternative to computing conservation scores using protein family position-specific scoring matrices (PSSMs)^46^, and allows approximating evolutionary fitness even in *de novo* designs with no homology information available. ESM-IF likelihood is a structure-conditioned score that quantifies how compatible a sequence is with a given backbone. Both ESM-1v and ESM-IF sequence likelihoods were shown to correlate with protein stability, activity, and expression^47,48^.

ProteinQC was applied to all 361 accepted *de novo* designs combined from the three example tasks: oxidoreductase scaffold design, PD-1 interface scaffold design and insulin receptor binder design (Figure 5a). The distribution of sequence entropy reached lower compared to the PDB reference set (Figure 5a top panel, left), closer inspection revealed sequences containing Lys/Glu repeats or stretches of Ala. The ESM-1v likelihood distribution for the 361 designs fell within the range of the distribution of the reference PDB set, albeit at the lower end of the tail (Figure 5a, top panel, middle). This indicates that *de novo* designs are generally well within the “fit” areas of the sequence space as considered by protein language models trained only on natural sequences. Although they may slightly deviate from the statistical patterns learned from natural protein sequences, presumably due to design constraints, reduced diversity, and absence of evolutionary context, scoring lower on the fitness spectrum might be explained by bias due to memorization of natural protein sequences found in the PDB reference distribution. On the contrary, the distribution of ESM-IF likelihood was placed right in the center of the reference PDB distribution (Figure 5a top panel, right). This is somewhat expected, given the designs were generated using ProteinMPNN which is a similar structure-aware design method.

**Figure 5.**
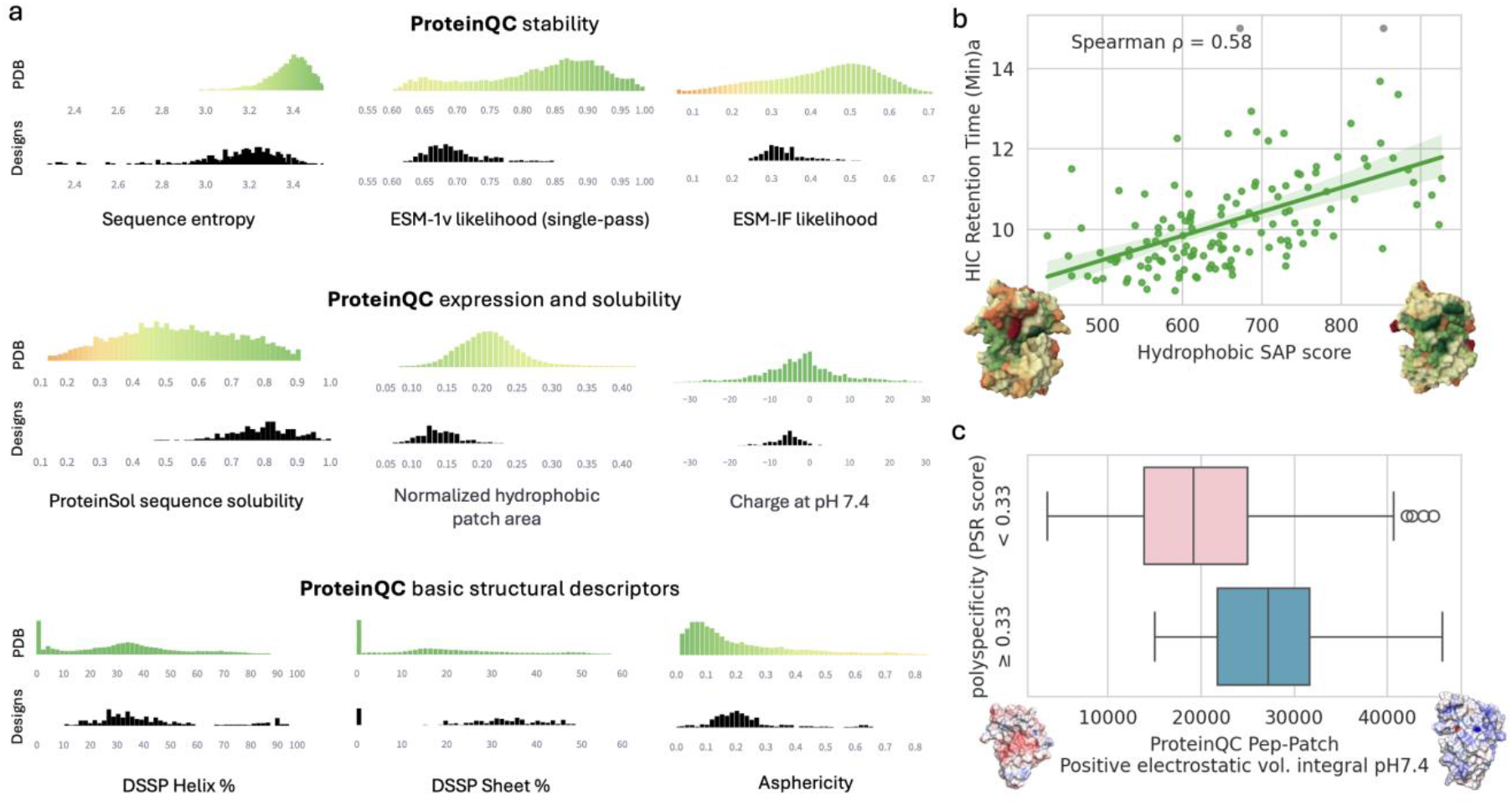
ProteinQC analysis. (a) Notable ProteinQC descriptors related to stability, expression, solubility, and basic structural properties for all 361 accepted de novo designs produced by the three example tasks: oxidoreductase scaffold design, PD-1 interface scaffold design and insulin receptor binder design. Top histograms show reference (full PDB) distributions colored using custom green-yellow-orange color scales corresponding to optimal, neutral, and sub-optimal regions of the distribution respectively; bottom histograms show observed distributions for designs of interest (b) Applicability of ProteinQC on antibody lead selection and engineering – HIC hydrophobicity assay of therapeutic antibodies from Jain dataset correlates with hydrophobic surface patch properties as computed by PEP-Patch SAP score. Example low-hydrophobicity and high-hydrophobicity structure indicate hydrophilic (red) and hydrophobic (green) regions using Mol* (c) Polyspecificity of antibodies of human mAbs from Shehata dataset correlates with positive surface charge as computed by PEP-Patch positive electrostatic volume integral. Example low positive patch content and high positive patch content structures indicate negatively charged (red) and positively charged (blue) patches visualized using PyMol APBS.69,70

#### De novo designs display expression and solubility patterns similar to well-behaved natural proteins

Ovo’s ProteinQC workflow computes multiple descriptors that evaluate designs’ solubility and expression liabilities (Supplementary Table S1). Three descriptors stand out. First, a scaled ProteinSol score (0–1) estimates solubility from amino-acid composition, with value >0.45 indicating above average solubility. Second, a normalized hydrophobic patch surface fraction (scaled 0-1) that measures the fraction of solvent-exposed surface occupied by hydrophobic patches, with elevated values indicating higher aggregation and insolubility risk. Third, net charge at pH 7.4 flags extreme or uneven charge distributions that can reduce solubility, promote non-specific interactions, or affect downstream developability. The distributions of these descriptors for the 361 *de novo* designs showed overall improved sequence-based solubility, lower fractions of hydrophobic-patch surface area, and slightly lower net charge at pH 7.4. Collectively, these results indicate that expression and solubility patterns are comparable to those observed in natural proteins (Figure 5a, middle panel).

Ovo’s ProteinQC workflow was also applied to a pool of uploaded clinical-stage monoclonal antibody (mAbs) structures with experimentally determined pre-developability assay measurements^49^, revealing a good correlation between Hydrophobic SAP score and HIC retention time (hydrophobic interaction chromatography, Spearman’s ρ = 0.58, Figure 5b). HIC retention increases with exposed hydrophobicity, since proteins with larger hydrophobic patches bind more strongly to the nonpolar HIC resin and elute later. A correlation between mAbs polyspecifity reagent assay (PSR)^50^ and PEP-Patch’s positive electrostatic volume integral at pH 7.4 was also apparent (Figure 5c). This descriptor quantifies the extent and magnitude of exposed positive patches. Large positively charged surface regions on antibodies can increase polyspecificity since they promote non-specific electrostatic interactions with a wide range of negatively charged molecules in the cell (e.g. acidic proteins, nucleic acids etc.) or in binding assays such as PSR.

#### Structure-related descriptors may shed light on packing, stability, and aggregation propensity

Structure-related descriptors capture a protein’s secondary-structure composition and overall three-dimensional shape calculated using DSSP and custom Python logic (Supplementary Table S1). Notable examples include the helix percentage, sheet percentage, radius of gyration, and asphericity. These features help distinguish helix-rich from sheet-rich folds, inform on folding and aggregation tendencies, and allow screening for compact, globular structures that are more likely to be stable and soluble. The distributions of these structure-related descriptors for the 361 *de novo* designs revealed comparable helix and sheet percentage to the PDB reference, indicating adequate degree of complex long-range beta topologies across design (Figure 5a, bottom panel, middle). Similar pattern was observed for the asphericity descriptor in the context of reference PDB distribution (Figure 5a, bottom panel, right), implying that the analyzed designs have a similar compactness and globularity profile to these of natural proteins, but also revealing a long tail of aspherical designs. Upon further inspection, these revealed elongated helical bundles that could be filtered out by the user depending on the design task. ProteinQC descriptor values for the 361 designs are provided in Supplementary Table S3.

Although we highlighted only a few descriptors here, the full descriptor set provides complementary, interpretable signals of sequence plausibility, predicted stability, surface aggregation risk, and geometric plausibility. Together they let users contextualize, cluster, and filter large design pools and prioritize candidates with balanced stability, expressibility, and foldability for experimental follow-up.

### Ovo plugins - a call to the community to develop models and tools in a shared ecosystem

Ovo plugins are lightweight Python packages that let developers and users rapidly integrate new tools and pipelines into a shared ecosystem. To maximize portability, reproducibility, and scalability, Ovo leverages the mature Nextflow language so pipelines can be constructed from reusable modules and run on any compute environment. A plugin can be a complete Nextflow pipeline, a modified variant of a pipeline providing an alternative method for a given design step, a minimal Nextflow pipeline wrapping a single design or analysis tool, or a visualization interface on top of the Ovo database. Plugins are intentionally easy to build. Once a container is defined (typically <50 lines of code) and a minimal Nextflow pipeline is implemented (<100 lines of code for simple methods), a simple user interface can be implemented in a single Python script (a few hundred lines of code), enabling authors to publish methods quickly and allowing users to adopt those methods with minimal integration overhead. Each plugin can be developed independently and published on GitHub or Python Package Index (PyPI), allowing installation using a single pip install command.

To demonstrate the versatility of Ovo, the power of its plugin architecture, and the ease of plugin development, we integrated ProteinDJ,^51^ an end-to-end Nextflow pipeline for *de novo* protein design described in a recently published preprint. To illustrate the straightforward integration of protein analysis tools, we implemented an example Ovo plugin for Promb^*^, a humanness evaluation toolkit based on average number of mutations to nearest peptide in a reference proteome.

We therefore call on model, method, pipeline, and user-interface developers to publish plugins for Ovo, a community-shared ecosystem. This will ensure a steady flow of unified, validated, and standardized pipelines so the community can share, reproduce, and scale high-quality design workflows.

## Discussion

The recent protein design revolution has produced an overwhelming proliferation of models, tools, and pipelines that have driven remarkable experimental successes in *de novo* design of peptides, mini-protein binders, enzymes, and antibodies, transforming the future of therapeutics, diagnostics, and manufacturing. At the same time, the sheer pace and diversity of development have made it difficult to choose, implement, test, integrate, deploy and execute experimentally validated pipelines. This is largely because tools are typically distributed as standalone code, notebooks, or ad hoc scripts, making installation, integration, data management, and compute provisioning non-trivial. Likewise, stitching multiple models into scalable design and analysis pipelines requires both deep domain expertise and substantial engineering effort. As a result, even promising design workflows may face slow adoption and limited community-wide benchmarking. Ovo was developed to address these challenges, not only providing a one-stop, extensible ecosystem that lowers the barriers to integrating, deploying, running, and evaluating *de novo* protein-design workflows, but also enabling efficient design and triage at industrial scale and speed.

### Ovo offers a state-of-the-art ecosystem for protein design

Interestingly, during preparation of this manuscript, ProteinDJ^51^, a protein design Nextflow pipeline, was released, underscoring the growing need for modular, scalable tooling in the field. The RFdiffusion pipeline in Ovo and ProteinDJ share the same core design philosophy. Both are Nextflow-based, both parallelize via batching, and both implement the standard end-to-end workflow (i.e. RFdiffusion, ProteinMPNN, optional FastRelax, AlphaFold2 validation). Both ProteinDJ and Ovo provide alternative sequence design methods, ProteinDJ providing FAMPNN^52^ while Ovo provides LigandMPNN building on OpenFold^16,53^ side-chain packing, so the workflow can run without a PyRosetta dependency. Regarding key differences between the two pipelines, Ovo implements predicted structure validation using the ColabDesign library^15^, which supports both AlphaFold2 monomer and multimer models and advanced template inputs, and additionally incorporates ESMFold. Ovo assigns stable unique identifiers to every design, preventing cross-run confusion and enabling cleaner result tracking via its robust storage layer. ProteinDJ on the other hand provides native support of additional protein design use cases such as symmetric scaffolding, fold-conditioned binder design, and enables automated exploration of input parameters. To illustrate the extensibility of Ovo, we integrated the ProteinDJ pipeline as a plugin, which attests to how quickly community tools can be combined to expand the shared Ovo ecosystem. Similar efforts to centralize protein engineering tools, such as TRILL^54^, are valuable and further emphasize the need for an ecosystem. However, they rely on a single Python environment and therefore inherit several sustainability and extensibility limitations. Regarding Ovo’s ProteinQC module, Stam and Wood developed DE-STRESS (DEsigned STRucture Evaluation ServiceS)^55^ which builds on a similar idea of computing protein descriptors such as secondary structure or hydrophobic patch areas and comparing them to a reference distribution such as the PDB. Compared to the ProteinQC module, DE-STRESS allows contextualizing against user-uploaded reference sets. However, unlike ProteinQC, it relies on tools that require commercial licenses, creating barriers to broader adoption.

### From limitations to opportunities: the future of Ovo

Ovo is a powerful platform, but it is important to note its limitations. Although Ovo can run on a local machine, many underlying methods are compute-intensive, therefore a heavy duty HPC or cloud infrastructure is required to perform designs at scale and unlock the platform’s full potential. *De novo* protein design remains research-intensive, so significant domain and computational expertise is still needed to internalize, implement, execute, and evaluate new models and tools. Likewise, there is an inherent trade-off between standardization and customization where standardized, reusable pipelines improve reproducibility and throughput but cannot cover every bespoke design need. Integrating third-party workflows can be challenging because converting external pipelines into Nextflow syntax may require substantial refactoring. To ease this burden, Ovo provides a programmatic Python API access and a results database so teams can run design workflows reproducibly from Jupyter notebooks, store outputs in Ovo database, browse them in the UI, and leverage Ovo for downstream evaluation and filtering. The plugins architecture provides straightforward extension points, and modalities not yet supported in the current release such as nanobodies and antibodies are prime candidates for community extensions. There are additional opportunities to establish community best practices and pre-defined filtering and thresholds. For example, the AlphaFold2 or AlphaFold3 derived interaction prediction score from aligned errors (ipSAE)^56^ might be a good candidate as the preferred metric for binder evaluation versus other metrics. Similar opportunities include expansion of the ProteinQC module to include more descriptors and implementation of additional clustering approaches and visualization options. ProteinQC currently supports reference sets drawn from subsets or the entire PDB. Thus, extending Ovo to support bespoke reference collections, for example THPdb databases^57,58^ of FDA approved therapeutic peptides and proteins, may better contextualize designs for specific applications. We therefore invite the community to contribute plugins, extensions, metrics, and workflows via the Ovo GitHub repository, which will serve as the central place to track work, surface issues, and review and merge contributions. By developing tools in a unified fashion, benchmarking and setting standards, the community can accelerate protein design across therapeutics and beyond.

## Methods

### Application architecture

Ovo is a Python application composed of multiple modules. The core module encapsulates all the functional logic components. Persistence is handled through two mechanisms: Ovo Storage and Ovo Database. Ovo Storage is a generalized interface for storing files such as PDB structures. It currently supports local filesystems, shared filesystems, or AWS S3 storage. Ovo Database uses a simple but robust data model to store designs, descriptors, jobs, and their metadata, as described in the next section. By default, Ovo uses SQLite, a single-file database, but supports any SQL database backend through SQLAlchemy. Ovo Scheduler is a general interface that provides functions for workflow submission, job status monitoring, and job output synchronization. Ovo currently provides two Scheduler implementations: 1) Nextflow Scheduler, which supports local execution as well as other Nextflow-supported platforms such as SLURM or PBS HPC executors and AWS Batch cloud executor, and 2) AWS HealthOmics Scheduler, which enables execution of Nextflow, WDL, or CWL pipelines using the AWS HealthOmics platform. Ovo provides a command-line interface that supports ad-hoc submission of workflows and individual tools. The Ovo Python codebase is organized such that the core logic functions can be imported from the Ovo Python module, providing access to designs and descriptors to enable their downstream analysis and visualization in Jupyter or other platform of choice.

### Database

In the Ovo data model, a Design corresponds to one designed molecule defined by its sequence. Designs are organized into Pools, where each Pool is created by a single submitted workflow. Submission parameters and job status metadata are stored in a separate DesignJob model using a workflow field storing a JSON-serialized instance of a DesignWorkflow dataclass. The DesignWorkflow dataclasses define workflow parameters and encapsulate use-case-specific logic. New use-cases can be added by Ovo plugins that implement their own DesignWorkflow subclasses. Each pool is identified by a randomly generated three-letter code (such as “buv”) unique within the current database instance. This ID is used as a prefix for the ID of each Design along with the backbone and sequence identifier (such as ovo_buv_0045_cycle01), ensuring that each design can be uniquely tracked to its job of origin (and the corresponding workflow parameters). Pools can be grouped into Rounds, corresponding to the iterative design-make-test-analyze loop, and further into Projects, enabling sustainable organization in a multi-user setting. Each design can be annotated with descriptors using the DescriptorValue model, which is a key-value “tall” representation where all descriptor values are stored in a single “value” column and associated with the descriptor using a “descriptor_key” column (as opposed to a “wide” representation which would use a separate column for each descriptor). Each DescriptorValue is associated with a single Design, a descriptor_key, and a DescriptorJob, storing workflow parameters in a DescriptorWorkflow dataclass, analogous to design jobs. Different types of Descriptors (such as AlphaFold2 pLDDT, Rosetta ddG, or total positive patch area) are uniquely defined by their descriptor_key and provide additional metadata such as their human-readable name and description. The Descriptor type objects are not defined in the database but inside the Ovo codebase and extended by plugins. Protein structures and other files are not stored directly in the database but referenced by their storage path as managed by the Storage class. The structure path string can be stored as a field of the Design model or as a special type of Descriptor. A diagram of the Ovo data model is provided in Supplementary Table S4.

### Pipeline definitions

Ovo leverages Nextflow^27^ to define standard design protocols as “pipelines” with defined input files and parameters, wrapping the end-to-end design process and producing the final designs as output files without need for user interaction in between. The nextflow.config file specifies the Nextflow input parameters and execution settings, while the main.nf script determines the main workflow as a dynamic directed acyclic graph (DAG) using custom groovy-based domain specific language (Nextflow DSL2). A Nextflow workflow is designed with a modular architecture where individual computational steps are encapsulated in containers or dedicated Conda environments to isolate each process’s dependencies and ensure reproducibility. For example, Ovo’s end-to-end RFdiffusion workflow entails multiple subsequent tasks for backbone and sequence generation, structure prediction validation, and filtering steps with conditional blocks. Pipeline execution starts with submitting a workflow with user-specified parameters, either through the Ovo CLI or via UI.

In case of Nextflow scheduler, a *nextflow run* subprocess is executed in the background, automatically orchestrating grid job submission and monitoring, enabling parallel execution of tasks and automatic retries. When this job is submitted via the web UI, job status is periodically checked and updated in the Ovo Database under a DesignJob object. Upon workflow completion, the results need to be “processed” by Ovo logic specific to the given pipeline, including copying relevant files from the workflow output directory (“publishDir” in Nextflow terminology) to Ovo Storage, assigning unique Design IDs and creating Design and DescriptorValue entries in the database.

### User installation and setup

The only required dependencies for Ovo are Python, Java 17 or later, and a container or environment management system such as Conda, Docker, Singularity, or Apptainer. Ovo can be installed via pip, which automatically installs Nextflow as a dependency. Upon first use, the ovo init command can be used to initialize the Ovo “home directory”, creating the necessary folder structure (workdir, storage, reference_files), default configuration file (config.yml) and SQLite database (ovo.db). The config.yml file contains settings for Ovo such as database connection or definitions of multiple scheduler instances, each with its own configuration to support different execution environments (e.g., local, HPC, or cloud) or execution settings (GPU, CPU-only, queue settings, etc.). Nextflow manages containerization according to the configuration settings. During Ovo startup (either in CLI or in the web server), Ovo checks for the Ovo home directory configured for the current installation of Ovo (or defined using OVO_HOME environment variable) and uses it to customize the logic for the user. Multiple independent Ovo home directories can be initialized on one system with one or more installations of Ovo. When using Conda (default), environments are created automatically as needed during workflow execution. For increased reliability and reproducibility, use of containers is recommended. Containers for specific workflows need to be built manually using instructions found in Ovo repository, although prebuilt container images are planned for future releases. Reference files such as trained model weights required for pipeline execution must be downloaded manually into the Ovo home directory. Common reference files, however, can be automatically retrieved using the ovo init command-line interface, simplifying the setup process.

### Web application

The Ovo web application is implemented using Streamlit, a Python framework that allows for quick implementation of interactive data-centric applications. The web interface enables workflow submission through step-by-step sections that interactively guide the user through all required input parameters customized for each use-case. Structures of workflow inputs, selected functional motifs, and generated designs are visualized interactively using Mol* structure viewer^59^. New pages can be added to the web interface via separately installable plugins. The Ovo web application can be executed locally, or in a multi-user setting on an institution’s server or HPC node or using platforms such as Posit (https://posit.co).

### RFdiffusion scaffold design

The end-to-end scaffold design pipeline starts with RFdiffusion^11^ for backbone generation. User inputs a structure in PDB format and chooses the functional motif residues to maintain fixed and the lengths of the redesigned segments by specifying a standard RFdiffusion “contigs” format. When submitting from CLI or using Python API, multiple contigs are allowed at once, for example supporting generation of different lengths of generated segments or permutations of fixed segments. By default, RFdiffusion runs 50 denoising steps and weights are set to “base”. To preserve the geometry of a small functional motif, it is preferable to choose the “active site” weights. After RFdiffusion backbone generation, backbone metrics are calculated to allow for preemptive design filtering, if selected by the user. Then, sequence generation is performed using LigandMPNN with ProteinMPNN model weights and side-chain packing enabled (one pack per design) to predict side-chain rotamers. Default temperature is set to 0.1. For *in silico* validation and filtering, AlphaFold2 initial guess is used by default, as done in the ColabDesign pipeline^15^. The inputs to AlphaFold2 are initialized with the coordinates of the designed structure from LigandMPNN output, followed by 3 “recycles” of AlphaFold2 prediction using model_1_ptm weights. By default, fixed functional motif regions that come from the original input structure are provided as structure template input. Template input can be disabled by the user in advanced settings to provide a more unbiased prediction. For monomer scaffold design, users can also validate using ESMFold^4^. The generation and evaluation process is divided into batches, each batch (100 backbones by default) is composed of a series of backbone generation, sequence design and structure prediction. Such batches can then be executed in parallel, enabling high throughput and short turnover time (provided that sufficient hardware resources are available).

For oxidoreductase motif scaffolding, PDB 1A4I chain A residues 56, 100 and 125 were used as fixed input motif, generating 10 to 100 residues in between and on N/C termini, with a restriction of desired length of exactly 150 amino acids. In total, 100 backbones * 6 permutations (of the order of the three fixed residues) * 8 sequence designs were generated, totaling 4800 designed sequences which were filtered with AlphaFold2 (PAE < 5, pLDDT > 80, Design RMSD < 2 Å and Native Motif RMSD < 1.5 Å), resulting in 13 accepted designs when RFdiffusion active site weights were employed. Running the same workflow with default RFdiffusion weights yielded no accepted designs due to higher Native Motif RMSD.

For PD-1 interface scaffolding, PDB 5IUS chain A segments 119-140 and 63-82 were joined with 15 to 40 generated residues, with additional 0 to 30 residues generated on N/C termini. Residues facing away from the interface (chain A residues 63, 65, 67, 69, 71, 72, 76, 79, 80, 82, 119-123, 125, 127, 129, 130, 133, 135, 137, 138, 140) were masked to RFdiffusion using sequence inpainting and redesigned with ProteinMPNN to allow for better packing against the designed scaffold. In total, 1000 backbones * 5 sequence designs were generated, totaling 5000 designed sequences which were filtered with AlphaFold2 (PAE < 5, pLDDT > 80, Design RMSD < 2 Å and Native Motif RMSD < 2 Å), resulting in 275 accepted designs. Running the same workflow without inpainting yielded only 1 accepted design, although this was mostly due to higher native motif RMSD of the inward-facing residues where the sidechains are expected to move, so a less stringent threshold could be applied. Workflow parameters and summary statistics for both example design tasks are provided in Supplementary Tables S2 and S3.

### RFdiffusion binder design and diversification

The end-to-end binder design pipeline starts with RFdiffusion backbone generation. The output structures from RFdiffusion are standardized to contain binder as chain A (numbered consecutively from 1) and target chain(s) as chain B (numbered same as in the input PDB). Sequence generation is performed by multiple cycles of ProteinMPNN and Rosetta FastRelax protocol^19^ (or LigandMPNN in case of no PyRosetta license). This entails full iterations of sequence generation and binding interface relaxation. Each cycle produces a different sequence design (57% mean pairwise sequence identity across sequence design cycles of a single backbone on average in the 2,150 generated insulin receptor binder backbones reported here). Designs from the final cycle as well as intermediate cycles are preserved and proceed to the following steps, where Rosetta^44,60^ ddG and other interface metrics are calculated, following the DLbinder protocol^19^. The designed structures and sequences are the input of AlphaFold2 multimer with initial guess for *in silico* validation and filtering. By default, structure of the target is provided as template, to account for missing MSA input, and to improve RMSD calculation when aligning to the target. In case when AlphaFold2 prediction is unable to find acceptable designs, advanced refolding settings allow to use also the structure of the binder chain or the full complex as AlphaFold2 structure template input. This can increase the likelihood of agreement between the predicted and designed complex, though it introduces additional bias toward the designed structure.

The same pipeline is used for binder diversification, the only major difference being the contig specification and the number of partial diffusions timesteps. Diversification starts with RFdiffusion partial diffusion protocol^14,20^ where target is kept fixed and binder backbone is diversified by applying a desired amount of noise followed by denoising (20 iterations by default). Multiple input structures are allowed, one contig specification per input structure. Thus, multiple designs can be partially diffused and re-designed, leading to diversification and optimization of a selected design pool. Akin to binder design workflow, sequences are designed through ProteinMPNN + FastRelax and AlphaFold2 multimer initial guess is used for validation. By default, all residues of the binder are partially diffused and subsequently redesigned with ProteinMPNN, but users can also partially diffuse only specific residues, making this workflow suitable for protein binder engineering tasks.

The workflow was applied to design a miniprotein binder of 50 to 100 residues against the insulin receptor PDB 4ZXB chain E residues 6-155, targeting hotspot residues 64, 88 and 96. In total, 1000 backbones * 3 ProteinMPNN FastRelax steps (+ 1 initial ProteinMPNN design) were generated, totaling 4000 generated sequences which were filtered with AlphaFold2 (iPAE < 10, pLDDT > 80, Target-aligned Binder RMSD < 2 A) and Rosetta (ddG < −30), resulting in 5 accepted designs. The same workflow with beta sheet RFdiffusion model weights yielded 1 accepted design with 20% of the structure forming beta sheets. The same workflow with LigandMPNN (ProteinMPNN model weights, 4 designs per backbone) used instead of ProteinMPNN FastRelax yielded 3 accepted designs. Running partial diffusion (50 backbones * 2 fastrelax steps) on three most AF2 confident RFdiffusion designs demonstrated the applicability of partial diffusion on exploiting similar areas of structure space: 37 out of 450 designs were accepted by the default AlphaFold2 and Rosetta filters, corresponding to 8.2% *in silico* success rate compared to 0.075% success rate of the full diffusion protocol. Workflow parameters and summary statistics are provided in Supplementary Tables S2 and S3.

### BindCraft binder design

BindCraft^18^ uses AlphaFold multimer backpropagation to design binders, SolubleMPNN^45^ for non-interface sequence design, and AlphaFold monomer and PyRosetta for *in silico* validation and filtering. By default, all five Alphafold multimer models are leveraged during BindCraft execution to avoid introducing model biases. One design run encapsulates iterative internal design generation until acceptance criteria are met for a specified number of designs, or when a time limit is reached. Unlike RFdiffusion where the number of generated designs is deterministic and controlled explicitly by the user so that it can be split into batches, parallelization of BindCraft is achieved by setting a desired number of “replicas”. These are parallel instances of the same BindCraft process, each typically running on one GPU instance and terminating when finding the desired number of accepted designs or when time limit is reached. The workflow settings and filters can be customized before workflow submission, starting from different presets as defined in the BindCraft repository (default/betasheet, 3-stage/4-stage, etc.). Internally, BindCraft uses PyRosetta metrics to score and filter designs, stored as descriptors for further downstream characterization and selection. All designs accepted or rejected by the built-in filtering logic are processed and stored in Ovo’s database. Trajectories that are discarded during the design process are not stored but can be accessed from the workflow working directory.

The workflow was used to design a miniprotein binder of 50 to 100 residues against the insulin receptor PDB 4ZXB chain E residues 6-155, targeting hotspot residues 64, 88 and 96.

Running for 6 hours on a single A10G GPU yielded 13 18 designs accepted by default BindCraft AlphaFold2 and Rosetta filters and 95 designs rejected by AlphaFold2 filters. No trajectories were rejected during the design protocol which could be attributed to AlphaFold2’s preference of the selected target epitope. Workflow parameters and summary statistics are provided in Supplementary Tables S2 and S3.

### ProteinQC

The ProteinQC workflow is implemented as a Nextflow pipeline that orchestrates the execution of individual tools as processes. Input PDB files are processed in batches in parallel, either for a single chain or for user-defined chain combination. For multi-chain selections, a single score is produced for the corresponding substructure. The workflow outputs CSV files containing one entry per PDB file and columns corresponding to computed descriptors. A complete list of available descriptors is provided in Supplementary Table S1.

Sequence-based tools include ESM-1v^46^, a protein language model trained on large numbers of sequences used to assess likelihood of native amino acid sequences. Protein-Sol^61^ for protein solubility prediction and ProtParam (ProteinAnalysis) from Biopython^39^ for calculating physicochemical properties and composition summaries. Structure-based methods include, DSSP^37,62^ for secondary structure assignment, and ESM-IF^6^, an inverse-folding protein language model used to assess likelihood of amino acid sequences given structural context. In addition, MDAnalysis^41,42^ for computing global and backbone structural metrics, and PEP-Patch^63^ for protein surface patch analysis.

To support interpretation of descriptor values, ProteinQC provides reference distributions generated from the Protein Data Bank (PDB)^64^. By comparing *de novo* designed protein structures to these empirical distributions, users can assess whether their descriptor values are typical of natural proteins. We provide precomputed descriptor distributions for the PDB, and its subsets based on structure source: human only, non-human only, and mixed human plus non-human (e.g., human target with non-human antibody) and on sequence length: short (≤100), medium (101–300), long (301–1000), and very long (>1000). To compute these reference distributions, PDB sequences were de-duplicated and processed in batches. For certain Biopython analyses ‘X’ was removed from the sequences. For ProteinSol, only sequences composed of the 20 standard amino acids and with length ≥21 residues (ProteinSol’s minimum supported length) were used. For ESM-1v, only sequences shorter than 1022 were analyzed due to the model sequence length limit. Although the tools were executed on deduplicated sequences, descriptor distributions reported were scaled to correspond to the full set of sequences. DSSP, ESM-IF, and MDAnalysis descriptors were calculated on PDB structures on each chain separately. For each descriptor produced by the tools above, non-null values were collected and trimmed to the empirical 1st–99th percentile to remove extreme outliers. Precomputed histograms were generated with NumPy using 50 bins and density=True. The same x-axis range was used for PDB subset histograms. Warning and error thresholds are defined for a subset of descriptors. For example, amino acid composition thresholds were derived from natural abundances, with warning and error levels set at 1.5× and 2× the natural abundance, respectively^65^.

### Plugins

Ovo supports third-party Python packages as plugins, enabling developers to independently extend Ovo with a new protein design method, characterization method, or user interface. A plugin is defined as a Python package, typically named ovo_plugin_name, defined using the standard new Python package pyproject.toml file, containing *ovo*.*plugins* as one of its entry points. The entry point references the plugin configuration module and variable, typically *plugin = ovo_plugin_name:plugin*, where *plugin* is a dictionary containing four keys: *pages, design_views, modules* and *descriptors*. During Ovo startup, the Python entry point logic retrieves all Ovo plugins among all packages installed in the environment. For each plugin, Ovo imports the package and adds any new pages and design views to the user interface. Descriptor objects that declare new types of descriptors are collected and appended to the application’s descriptor registry. To include a new design or descriptor workflow, a plugin package needs to contain 1) a Python script defining the Streamlit user interface, 2) a new DesignWorkflow or DescriptorWorkflow subclass that defines the input parameters and provides submission and processing logic functions, 3) new Descriptor objects that define any new types of design annotations including numeric, categorical, or references to files stored in Ovo Storage, and 4) a Nextflow pipeline that defines a simple single-step process or complex end-to-end workflows, abstracting execution logic and isolating software dependencies. Adding additional Python packages as direct dependencies of the plugin package is discouraged to ensure long-term compatibility of Ovo dependencies. Nextflow pipelines can be declared directly as part of the plugin package in its pipeline’s module, or as separate git repositories (compatible with the nf-core standard) that can be referenced by the workflow submission logic by their git URL. The current implementation does not allow plugins to easily provide drop-in replacements of individual stages of an existing pipeline. However, based on the nf-core philosophy, plugins are welcome to reuse the building blocks (modules) of existing pipelines to create adjusted versions of a pipeline, for example by swapping the backbone generation, sequence design, or structure prediction method. Please refer to Ovo code repository for real-world examples of Ovo plugins for design or analysis.

## Data availability

All code for this publication is available in the following GitHub repository: https://github.com/MSDLLCpapers/ovo.

We also deployed a demo web application for demonstrating design visualization capabilities at https://ovo.dichlab.org. Documentation is available at https://ovo.dichlab.org/docs.

*De novo* design workflow results presented in this publication are available through reproducible Jupyter notebooks at: https://github.com/MSDLLCpapers/ovo-examples.

## Supplementary data

Supplementary Tables S1-S4 include descriptions of all Ovo descriptors, example design task statistics, ProteinQC descriptor results, and the Ovo data model.

## Funding

This work was supported by Merck Sharp & Dohme LLC, a subsidiary of Merck & Co., Inc., Rahway, NJ, USA.

## Conflict of interest

All authors that are/were employees of Merck Sharp & Dohme LLC, a subsidiary of Merck & Co., Inc., Rahway, NJ, USA may hold stocks and/or stock options in Merck & Co., Inc., Rahway, NJ, USA.

## Acknowledgments

We thank Monika Pulcova, Viktorie Malickova and Patricia Sotakova for help implementing some supporting features, and Michal Drda for leading the product analysis efforts at the start of the project. We are immensely grateful to Dave Pace, Carol A. Rohl, Matt Studney for supporting this work and Aaron Fridman, Shahar Bar and Girish BhamBhani for nurturing the partnership between AWS and Merck & Co., Inc., Rahway, NJ, USA. We are grateful to Christopher Warren for his insightful guidance throughout the tool’s development, and to Thomas W. Linsky and Benjamin Jagger for their thorough review of the manuscript.

## Author Contributions

DB conceived the study. DP designed the solution. DP, TC, MA, AK, LP, and HH developed and implemented the entire application. DB and DP supervised the study. ND implemented the first version of the Nextflow pipelines and integrated with AWS HealthOmics services. DP and MA implemented the final version of the Nextflow pipelines, and TC implemented the ProteinQC pipeline. DB wrote the manuscript. AK, DP, MA and TC wrote the method section and contributed to the final draft. All authors reviewed and approved the final draft.

## Footnotes

* https://github.com/MSDLLCpapers/promb

## Notes

https://github.com/MSDLLCpapers/ovo

